# Sex-biased gene expression in *Drosophila melanogaster* is constrained by ontogeny and genetic architecture

**DOI:** 10.1101/034728

**Authors:** FC Ingleby, CL Webster, TM Pennell, I Flis, EH Morrow

## Abstract

Sexual dimorphism is predicted to be constrained by the underlying genetic architecture shared between the sexes and through ontogeny, but whole-transcriptome data for both sexes across genotypes and developmental stages are lacking. Within a quantitative genetic framework, we sequenced RNA from *Drosophila melanogaster* at different developmental stages to examine sex-biased gene expression and how selection acts upon it. We found evidence that gene expression is constrained by both univariate and multivariate shared genetic variation between genes, sexes and developmental stages, but may be resolved by differential splicing. These results provide a comprehensive picture of how conflict over sexual dimorphism varies through development and clarifies the conditions under which it is predicted to evolve.

---

Sexual dimorphism is widespread across animals and plants, and is likely to have evolved in response to the different reproductive roles of the sexes within a species^1^. However, in most species, the majority of genes are shared between males and females, and sex-specific fitness optima for shared traits can create conflict, called intralocus sexual conflict^2^. Intralocus sexual conflict is driven by sexually antagonistic selection, which pushes the phenotypic value of the trait in different directions. It is therefore thought that sexually antagonistic selection leads to the evolution of sexual dimorphism and ultimately the resolution of conflict, by allowing each sex to express the trait independently and to each achieve their optimum phenotype^3-4^.

For sexually dimorphic phenotypes to develop from the same genes, we might expect differences in gene expression (or in downstream modifications and regulation) between the sexes at some stage of development^5-6^. Sexually dimorphic gene expression, or sex-biased gene expression, has been examined in detail in a wide-range of species^7^. But, although it is known that there can be substantial variation in gene expression between developmental stages, less is understood about the sex-specific dynamics of gene expression throughout development^6-8-9^. This is a potentially significant gap in our knowledge in the face of evidence that gene expression during early development can strongly impact on adult phenotype^10-11^, as can the early environment more generally^12-15^

In terms of understanding how sex-biased gene expression relates to sexual conflict and conflict resolution, very few studies have directly associated sex differences in gene expression with sex-specific fitness^16-17^. Furthermore, our understanding of how the transcriptome is related to fitness is largely based on adult gene expression, and little is known about how sex differences laid down during development might influence overall fitness. Further research is necessary to synthesise a comprehensive picture of how sex-biased gene expression might mediate sexual conflict. We argue that a quantitative genetic and developmental perspective will provide valuable insights into the genetics and ontogeny of conflict and resolution, by enabling genetic covariance across sexes and developmental stages to be quantified and by testing the association of these patterns of gene expression with fitness.

Here, we present the first quantification of a transcriptome across sexes, developmental stages and genotypes, and interpret the variation in gene expression in terms of sex-specific fitness. We demonstrate widespread sexbiased gene expression in *Drosophila melanogaster* larvae, pupae and adults, as well as considerable differences in the fitness consequences of gene expression throughout development. We combine this developmental perspective with a quantitative genomic approach that has been used increasingly in recent research^16-19^. By analysing gene expression in different genetic lines, we find evidence for potential constraints on conflict resolution through genetically correlated gene expression between genes, across sexes and across development. Importantly, we use multivariate analyses alongside our univariate analyses, to account for genetic covariance between different genes, an aspect that is overlooked in more common univariate analyses of gene expression. In addition, although our data does not allow full examination of all potential downstream modifications to gene expression, we do explore a potential route for conflict resolution via differential splicing between males and females. In sum, our results offer new insight into the genetics of sex differences in gene expression throughout development, and how these differences could mediate sexual conflict

## Results

RNA was extracted from male and female third-instar larvae, pupae and adults from each of 10 hemiclonal lines, and gene expression was quantified by RNA- sequencing. Each hemiclonal line is generated through a series of crosses^17^ that result in each fly within a line (both males and females) sharing a haplotype that is expressed alongside a random haplotype from the base population, allowing additive genetic variation to be estimated for each trait. We examined the expression of 14008 genes in total, of which 13501,13602 and 13495 genes were expressed in larvae, pupae and adults respectively. Sex-specific fitness for each hemiclonal line (hereafter referred to as a ‘line’) was measured as reproductive success under competitive conditions.

### Sex-biased and sexually antagonistic gene expression

Initial analyses partitioned variance in the expression of each gene within each stage between sex, line and sex-by-line effects. Table 1A summarises the numbers of sex-biased genes at each stage (significant ‘sex’ term at FDR<0.001). Sex-biased expression was generally high throughout development (83.4% and 78.7% of larval and pupal genes, respectively), and highest in adults, with 89.9% of genes showing significant differences in expression between adult males and females. In larvae and pupae, sex-biased gene expression is predominantly male-biased, but in adults there are more equal numbers of female- and male-biased genes (Table 1A). Imposing the additional criteria of a fold change >2 reduces the numbers of genes called as sex-biased at each developmental stage, although this reduction is most noticeable in larvae and pupae, indicating that the magnitude of sex-biased expression is generally higher in adults (Table 1A).

**Table 1.** Overall numbers of genes that were (A) significantly sex-biased, (B) had significant between-line (genetic) and sex*line (sex-specific genetic) variation, and (C) significantly associated with sexually antagonistic (fitness*sex) or sexually concordant (fitness with no fitness*sex interaction) fitness. All results are taken from the linear models described in the text. In total, 13501,13602 and 13495 genes were tested for larvae, pupae and adults respectively. The percentages of these totals are shown italicised in brackets.

|  | **Larvae** | **Pupae** | **Adults** |
| --- | --- | --- | --- |
| **(A) Sex-biased** | 11262 (83.4) | 10705 (78.7) | 12133 (89.9) |
| **Sex-biased (fold change > 2)** | 4535 (33.6) | 2613 (19.2) | 8252 (61.2) |
| **Female-biased** | 1200 (8.9) | 3620 (26.6) | 5696 (42.2) |
| **Female-biased (fold change > 2)** | 131 (1.0) | 324 (2.4) | 3525 (26.1) |
| **Male-biased** | 10062 (74.5) | 7085 (52.1) | 6437 (47.7) |
| **Male-biased (fold change > 2)** | 4404 (32.6) | 2289 (16.8) | 4727 (35.1) |
| **(B) Genetic variation** | 2870 (21.3) | 4706 (34.6) | 4912 (36.4) |
| **Sex-specific genetic variation** | 192 (1.4) | 3110 (22.3) | 2043 (15.1) |
| **(C) Sexually antagonistic** | 841 (6.2) | 610 (4.5) | 1300 (9.6) |
| **Sexually concordant** | 420 (3.1) | 2493 (18.3) | 1659 (12.3) |

From the same models, significant genetic (between-line) and sex-specific genetic (sex*line) variation were identified at FDR<0.05 (Table 1B). The numbers of genes with significant genetic variation increased throughout development, from approximately a fifth of all genes tested in larvae, to just over a third of all genes tested in adults. Notably, sex-by-line variation - which indicates some extent of genetic variation for sex differences in gene expression - was very low in larvae (1.4%), but higher in adults (15.1%) and considerably higher in pupae (22.1%), suggesting a higher capacity for the evolution of sexually dimorphic gene expression in pupae. In a species with a holometabolous life cycle like *D. melanogaster*, the pupae represent the most dynamic phase of development in terms of tissue differentiation and the development of sex- specific phenotypes, whereas the larval stage mostly concerns growth rather than differentiation, regardless of sex. Although the samples were carefully staged, pupal metamorphosis is such a dynamic developmental stage that we cannot rule out the possibility that genetic variation for gene expression is confounded with genetic differences in the precise timing and process of metamorphosis. Nonetheless, the particularly high level of sex-specific genetic variation in pupae shows that we identified significant genetic variation for sex differences in pupal metamorphosis, consistent with our other findings.

Candidate sexually antagonistic genes were identified by regressing fitness against gene expression, where a significant sex*fitness interaction (FDR<0.05) was used to call candidate sexually antagonistic (SA) genes, and a significant fitness term in the model (FDR<0.05), with a non-significant sex*fitness interaction, interpreted here as sexually concordant (SC) fitness consequences (Table 1C). Overall numbers of fitness-associated genes (SA and SC combined) were highest in pupae, which consisted predominantly of SC genes. Numbers of SA candidate genes were highest in adults (9.6% of genes tested) and lowest in pupae (4.5%). The overlap in SA and SC calls between stages was low (Figure 1), suggesting that both forms of selection vary widely throughout development. For all tests of sex-biased expression, overall genetic variation, and fitness association, genes are identified in Supplementary File 1.

**Figure 1.**
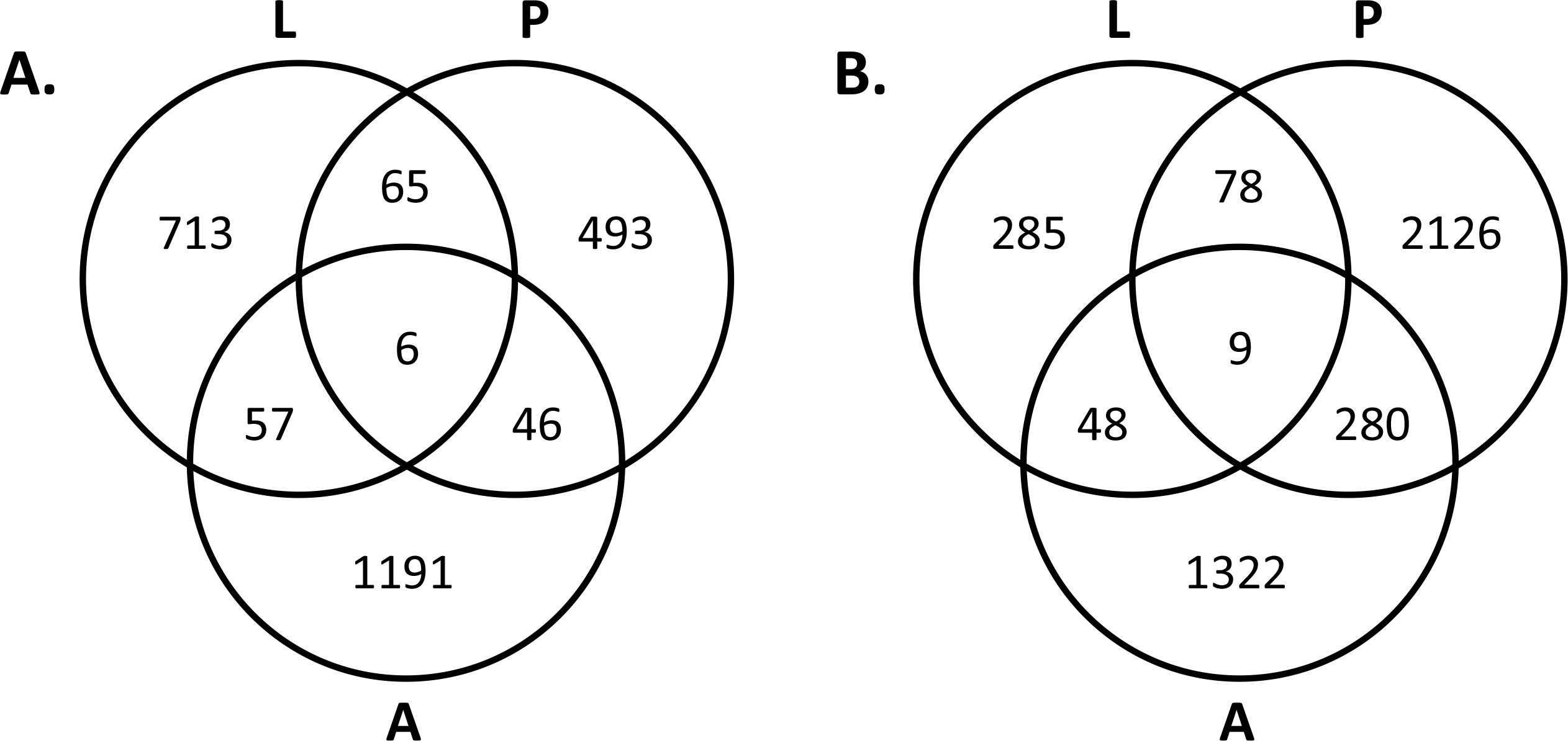
Venn diagrams of the overlap between (A) sexually antagonistic and (B) sexually concordant candidate genes called in larvae (L), pupae (P) and adults (A). In total, 2571 genes were SA in at least one stage (18.4% of the 14008 genes tested), and 4148 genes were SC in at least one stage (29.6%).

Across development, 2571 genes (18.4% of the total 14008 genes tested throughout development) were identified as SA candidate genes in at least one developmental stage, suggesting a higher overall level of conflict throughout the genome than estimated from analyses of adult gene expression alone in the same laboratory population over five years previously^17^. Notably, even within the adult subset of our data, there was low and non-significant overlap (approx.10%) between the genes that were called as SA candidates compared to the genes called as SA candidates previously^17^ Studies like these capture a snapshot of sexually antagonistic fitness variation at a given time point, but it appears that conflict and resolution are dynamic^17-20^.

### Modules of correlated gene expression

In an attempt to simplify a complex, whole-transcriptome dataset and identify patterns of potential functional interest, we identified modules of correlated gene expression among the fitness-related genes (SA and SC combined) within each developmental stage. First, we calculated genetic correlations for each pairwise combination of fitness-related genes for both sexes within each stage, and used the mean absolute value of the correlations for each gene with all other genes as a measure of gene ‘connectivity’^16^. Average connectivity was generally high, and highest in adults (Figure 2). In larvae and adults, connectivity was significantly higher for SA genes than for SC genes, but the opposite was true in pupae, although the absolute difference was small (Figure 2).

**Figure 2.**
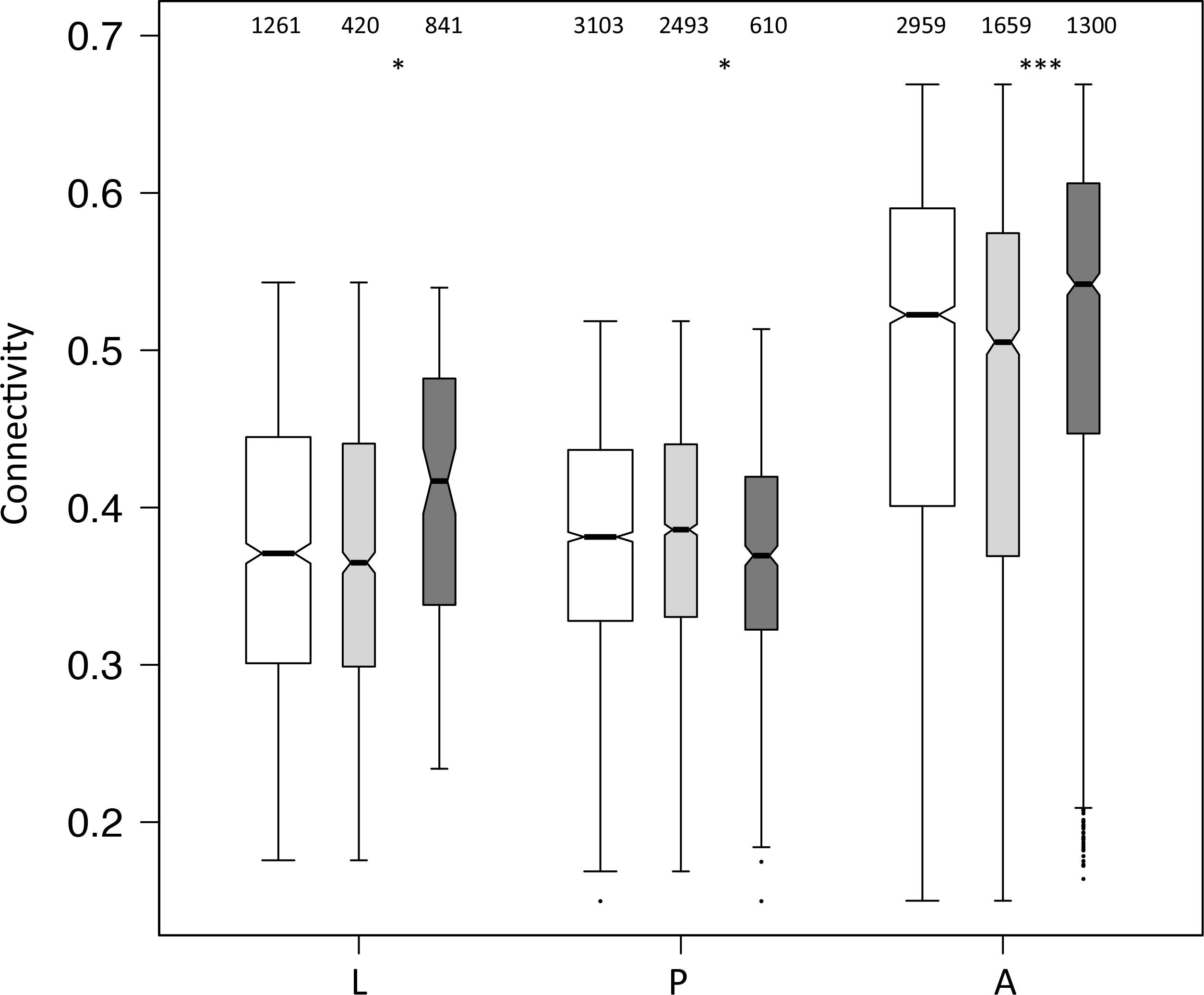
Connectivity of all fitness-related genes for each stage, as the average absolute genetic correlation of all pairwise gene combinations within each stage. White boxes represent the overall connectivity for all fitness-related genes in larvae (L), pupae (P) and adults (A). Smaller boxes represent connectivity of sexually antagonistic (dark grey) and sexually concordant (pale grey) genes separately. Numbers of genes included in each box are shown. Notches in boxplots represent 95% CI approximations, as ±1.58*IQR/√N. Asterisks indicate significance of the difference between SA/SC genes at each stage.

Next, we inferred functionality by using the pairwise correlations calculated above to cluster the fitness-associated genes into transcription modules within each stage^21^. We tested these modules for enrichment with SA candidate genes at each stage, and tested the largest modules (>100 genes) for enrichment with tissue-specific genes identified from the FlyAtlas database^22^ of larval and adult tissues (results in Supplementary File 2). There were 27 transcription modules identified in larvae, 5 of which were significantly enriched with larval SA candidate genes at FDR<0.05 (Figure 3A). A large larval transcription module (#11 in Figure 3A) was enriched for both SA candidate genes as well as salivary gland and malphigian tubule-specific genes, suggesting this module may be associated with feeding, and that larval feeding behaviour might have SA fitness consequences. In pupae, there were 56 transcription modules, of which only 4 were significantly enriched for pupal SA candidates (Figure 3B). The clustering results for pupal fitness-associated genes were dominated by one large transcription module (#36 in Figure 3B). This module, similarly to the largest transcription module in larvae (#27 in Figure 3A), was enriched for a variety of different tissue-specific genes, all from somatic rather than germ line tissues. In adults, we found 36 transcription modules, of which 8 were enriched with adult SA candidate genes (Figure 3C). Modules #5, 6 and 9 were heavily enriched with testes-specific genes, whereas modules #11 and 19 were enriched for multiple tissue-specific sets of genes, including a combination of head, brain and CNS tissue-specific genes, suggesting a putative link between behaviour and sex-specific fitness. These results are consistent with previous research that has demonstrated sexual conflict over adult locomotory behaviour in *D. melanogaster*^23^. Module #19 was also significantly enriched with adult SA candidate genes, and so these transcription modules may harbour interesting candidate genes for further research (Figure 3C).

**Figure 3.**
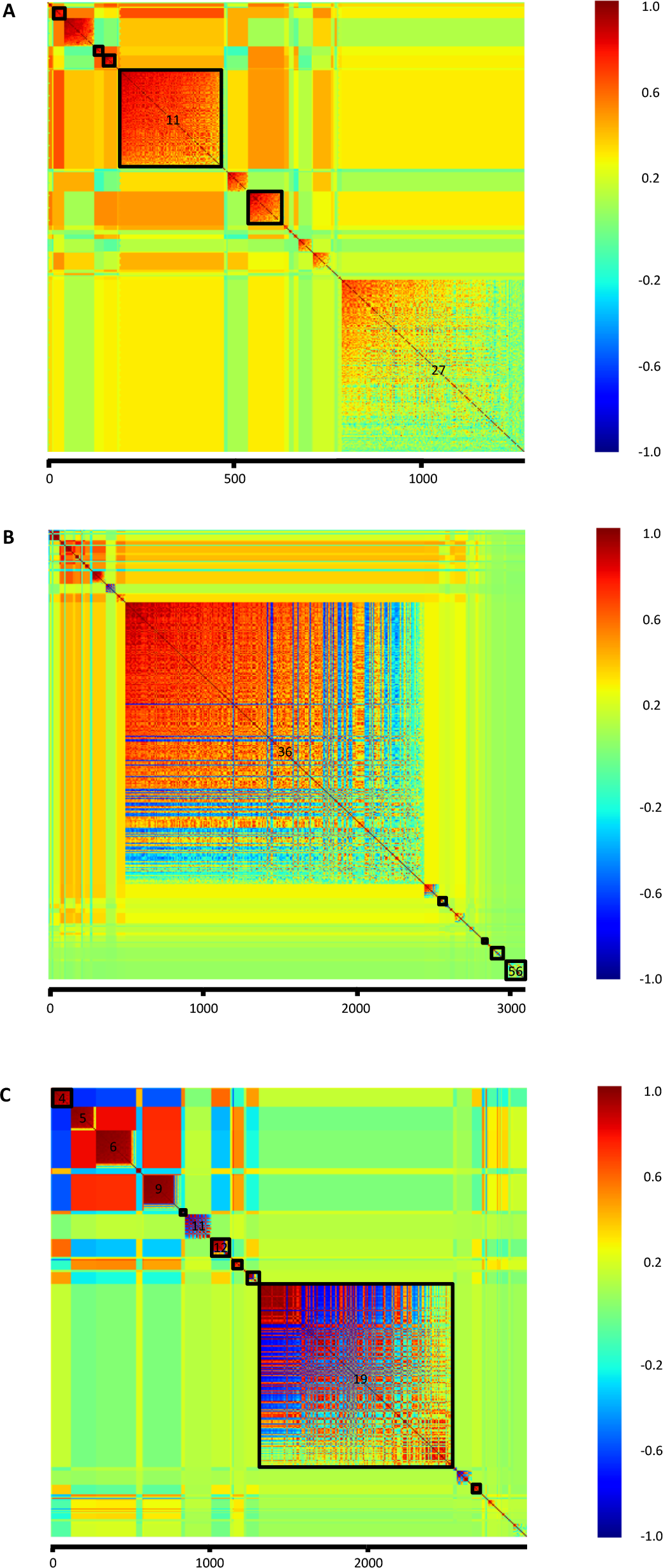
Modules of correlated gene expression for all fitness-related genes in (A) larvae (N=1261); (B) pupae (N=3103); and (C) adults (N=2959). Colours within modules represent genetic correlations between all pairwise combinations of genes; colours between modules represent the average genetic correlation between modules. Outlined modules are those that tested significant for enrichment with sexually antagonistic genes. Numbered modules are those with >100 genes that were tested for enrichment with tissue-specific genes.

### Shared quantitative genetic variation

To explore the potential for genetic constraints to hinder the resolution of sexual conflict over gene expression, we measured quantitative genetic variation underlying gene expression. In particular, we were interested in shared genetic variation between sexes and between developmental stages, as this shared genetic variation could prevent the independent evolution of sex- and stage- specific gene expression.

Initially, we considered genetic variation from a univariate perspective. For each gene with significant genetic variation in at least one developmental stage (N=8761), we ran a model that partitioned variance in gene expression between sex, stage and line, creating a 6x6 genetic variance-covariance matrix for each gene individually (2 sexes x 3 stages), as shown inset in Figures 4A-D. This matrix can be split into four sources of variance: (1) sex- and stage-specific genetic variance (Figure 4A), (2) between-sex genetic covariance within each stage (Figure 4B), (3) between-stage genetic covariance within each sex (Figure 4C), and (4) genetic covariance between both sexes and stages (Figure 4D). Covariance estimates were scaled to the total amount of genetic variation in the full matrix. Overall, genes that were identified as SA in at least one stage had significantly more sex- and stage-specific genetic variation than genes that were not SA in any stage of development (Figure 4A). This was expected, as theory predicts that SA selection will help maintain genetic variation^3-4^. There is also substantial shared genetic variation both between sexes and between stages, and interestingly this is significantly higher for SA genes than non-SA genes in all instances (Figures 4B-D). Notably, there is even evidence of considerable shared genetic variation between-sex and between-stage (Figure 4D), suggesting complex genetic covariance of gene expression across males and females at different developmental stages. The results of these models were also used to calculate the intersexual genetic correlation, r_mf_, for each gene at each stage. As might be expected from the results of overall between-sex genetic covariance shown in Figure 4B, r_mf_ is higher for SA genes than non-SA genes at each stage, although this difference is small in pupae (Figure 5).

**Figure 4.**
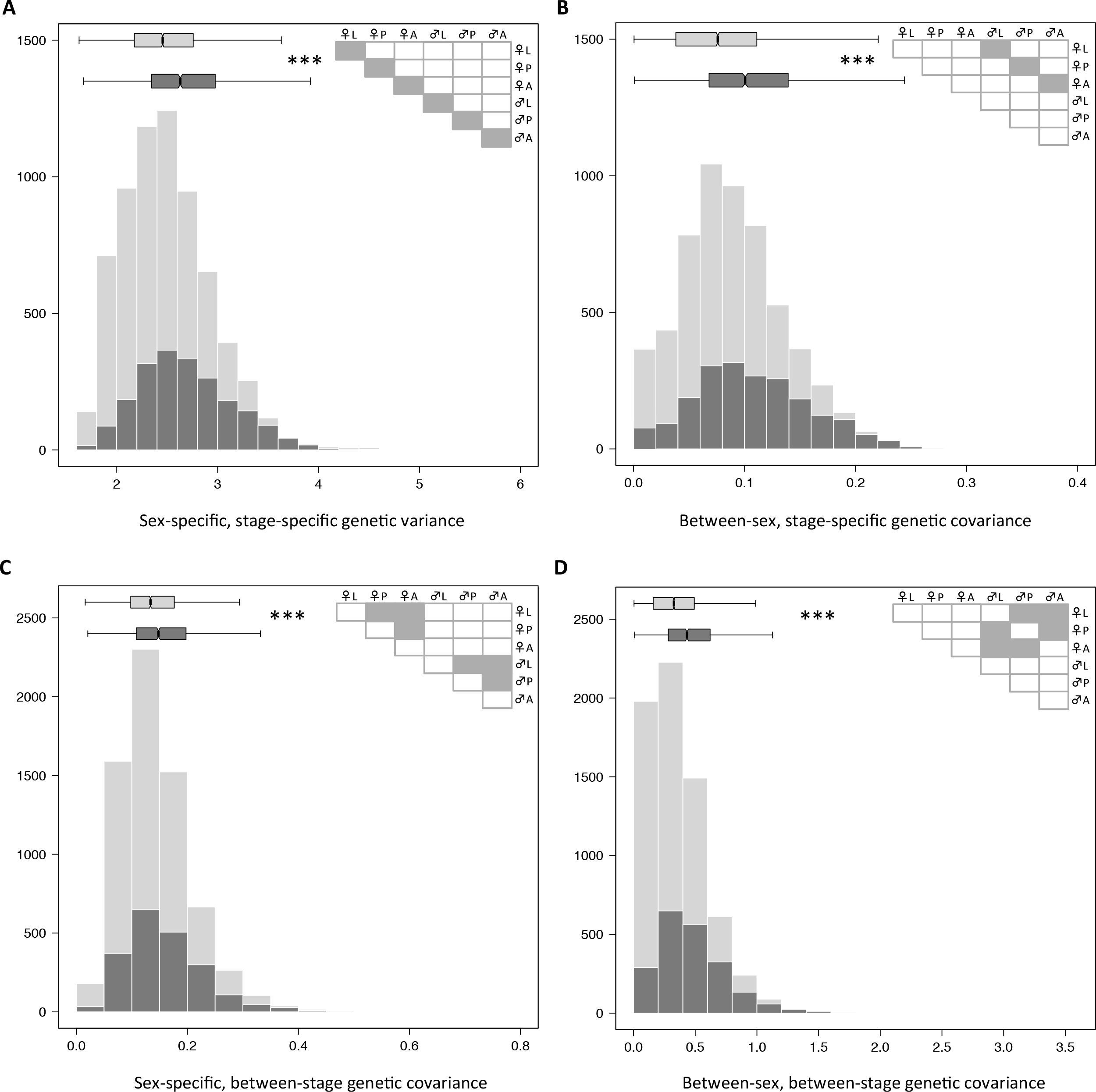
Histograms of the between-sex and between-stage genetic variance-covariance matrix components for individual genes, partitioned between variance components as shown in the inset matrix diagrams. (A) Sum of the sex-specific, stage-specific genetic variance. (B) Sum of the absolute values of the between-sex genetic covariance within each stage. (C) Sum of the absolute values of the between-stage genetic covariance within both sexes. (D) Sum of the absolute values of the between-sex, between-stage genetic covariance. Only genes with significant genetic variance for at least one developmental stage are included (N=8761). Genes that are sexually antagonistic in at least one stage (N=2056) are shown in dark grey and genes that are not SA at any stage are shown in pale grey (N=6705). Notches in boxplots represent ±1.58*IQR/VN. Asterisks indicate significance of the difference between SA/non-SA genes at each stage.

**Figure 5.**
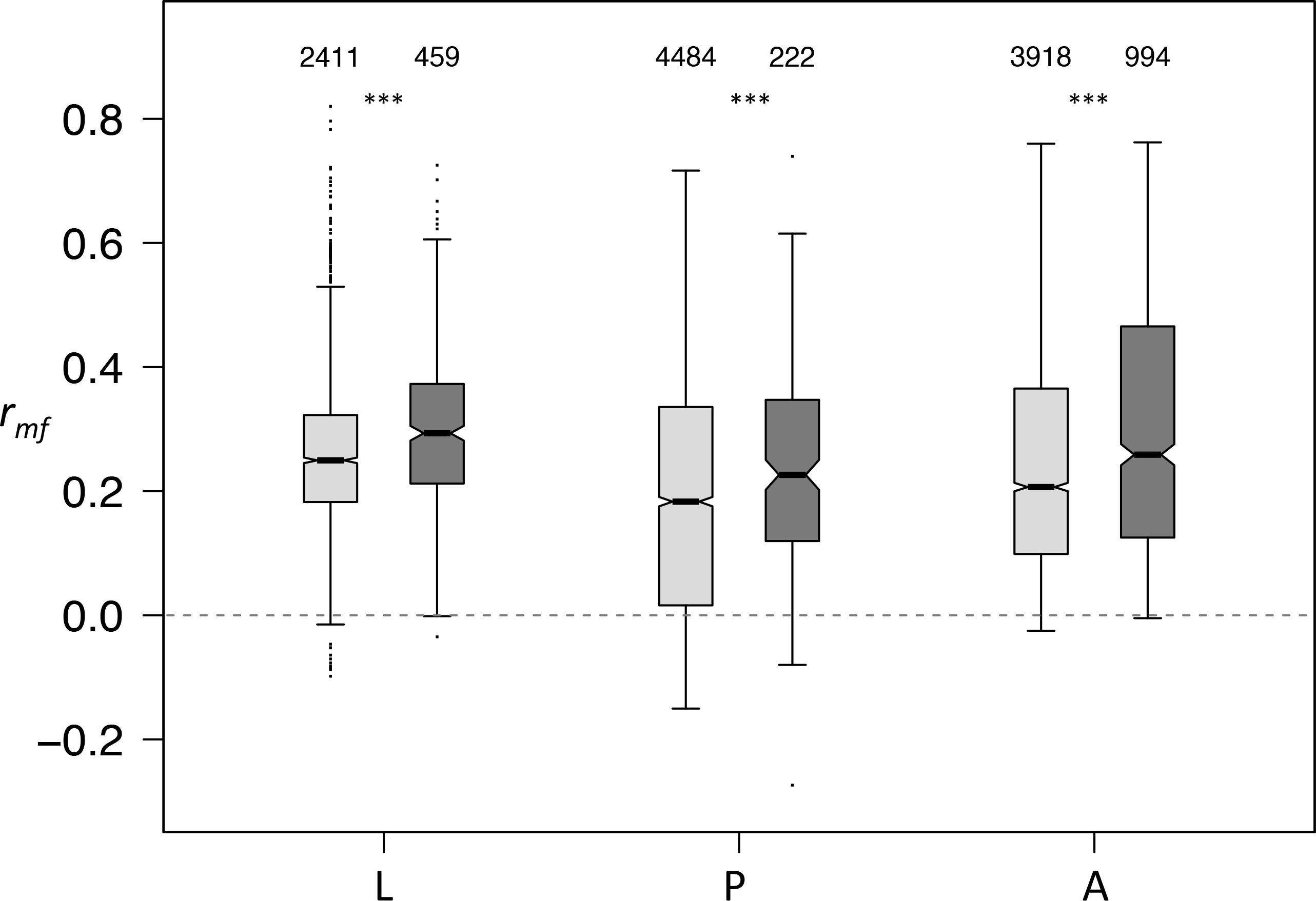
Intersexual genetic correlation (r_mf_) for sexually antagonistic (dark grey) and non-sexually antagonistic (pale grey) genes in larvae (L), pupae (P) and adults (A). Only genes with significant genetic variation at each stage are shown, and the numbers of genes included in each box are shown. Notches in boxplots represent ±1.58*IQR/√N. Asterisks indicate significance of the difference between SA/non-SA genes at each stage.

Next, we used multivariate analyses of quantitative genetic variation, combined with a re-sampling technique, to examine shared genetic variation between-sex and between-gene. This involved calculating the full genetic variance-covariance matrix for sub-samples of genes expressed in males and females **(G_mf_**, or **G** matrix). The **G** matrix includes the sub-matrix **B**, which summarises between-sex and between-gene genetic covariance^24^ (see Supplementary File 3). We estimated the average **G** matrix for SA and SC genes for larvae, pupae and adults independently. Overall, we found that **B** tended to have a higher magnitude for SA genes than for SC genes in larvae and adults, indicating that there is higher between-sex and between-gene genetic covariance in SA genes. The opposite was true for pupae (Figure 6A), consistent with the patterns of connectivity for SA genes compared with SC genes in Figure 2. We also estimated the matrix correlation between the upper and lower halves of the **B** matrix, and found that this was significantly higher for SA genes than for SC genes at each developmental stage (Figure 6B), although this difference was small in pupae. This shows that genetic covariances are more strongly correlated between sexes for SA than for SC genes, implying more potential for male and female gene expression to evolve independently in SC genes.

**Figure 6.**
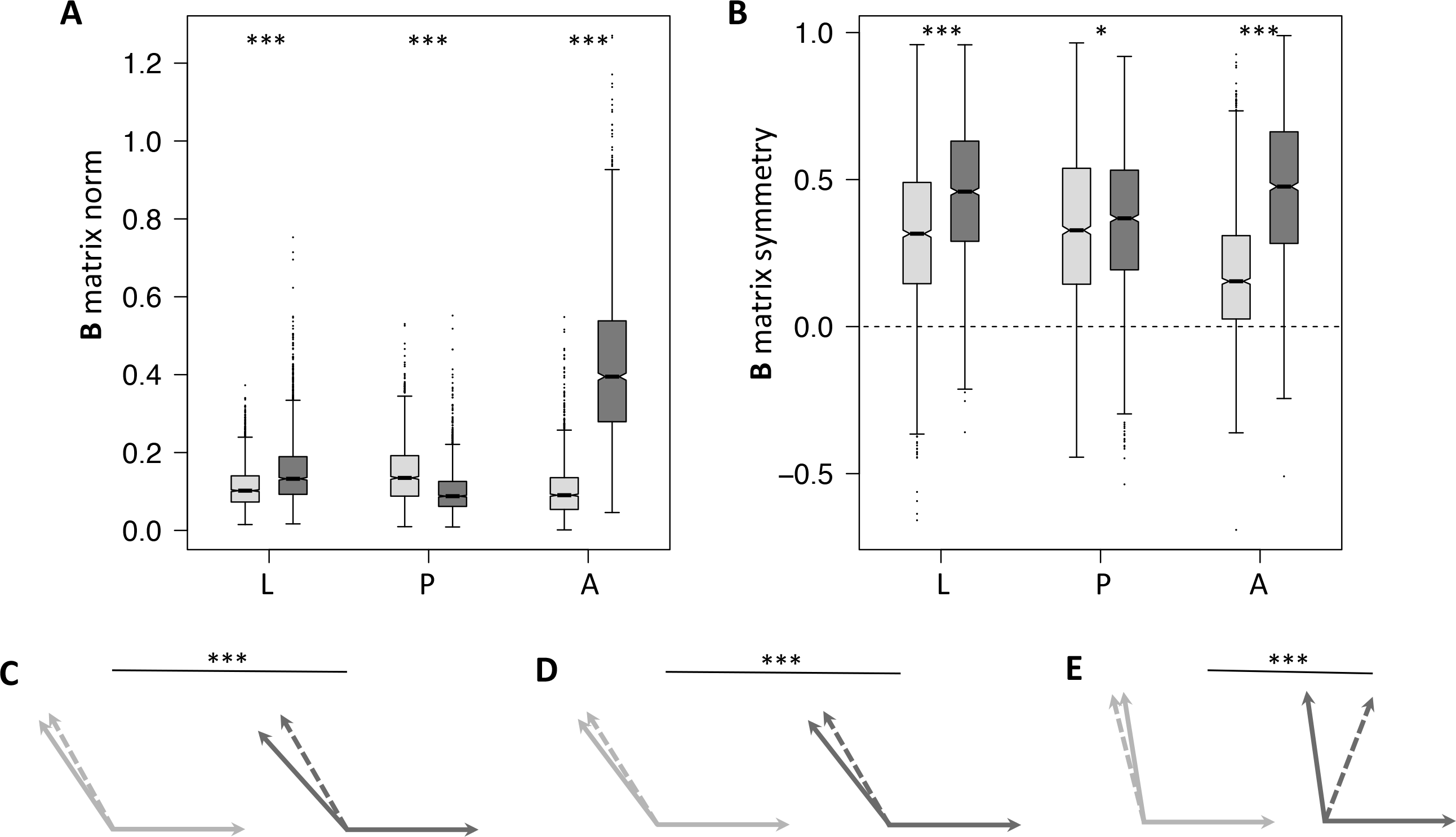
Genetic variation in the average **B** matrix (between-sex and between-gene) for SC and SA genes in larvae (L), pupae (P) and adults (A). (A) The **B** matrix norm, **∥B∥**, as a measure of the overall magnitude of genetic variation in **B** for SC (pale grey) and SA (dark grey) genes at each stage. Boxes represent the estimates from all 2000 iterations of the model, notches in boxplots represent ±1.58*IQR/√N, and asterisks indicate significant difference between SA/SC genes at each stage. (B) The symmetry of the **B** matrix, as the matrix correlation between the two halves of **B**, for SC (pale grey) and SA (dark grey) genes at each stage. Boxes represent the estimates from all 2000 iterations of the model, notches in boxplots represent ±1.58*IQR/√N, and asterisks indicate significance of the difference between SA/SC genes at each stage. (C-E) The angle between the male and female predicted response to selection without the **B** matrix (solid arrows) and adjusted for the inclusion of the **B** matrix (dashed arrow) for SC (pale grey) and SA (dark grey) genes in larvae (C), pupae (D) and adults (E).

Finally, we used the multivariate breeder’s equation^24^ to estimate the vectors of the predicted response to selection for males and females for SA and SC genes. We then use the angle between these vectors as a measure of the predicted divergence between male and female trait evolution. When the between-sex, between-gene shared genetic variation in **B** is included in these calculations, the divergence between the sexes tends to decrease, as shared genetic variation forces the predicted response to selection between the sexes to realign with one another to some extent. However, this realignment is stronger for SA than SC genes in larvae (Figure 6C), pupae (Figure 6D) and especially in adults (Figure 6E), suggesting that the multivariate between-sex genetic covariance within **B** presents a significant constraint on the independent evolution of the sexes, particularly in adults.

### Evidence for differential splicing

Differential splicing was examined by comparing splicing between male and female samples within each developmental stage, for all genes with evidence of alternative isoform expression. There was most evidence of differential splicing between the sexes in adults, where 1089 genes out of 2822 genes tested (38.6%) showed significant differential splicing between sexes (FDR<0.05). Evidence for significant sex-specific splicing in larvae (0.9%, 14/1636) and pupae (3.8%, 77/2015) was low. However, the overall extent of differential splicing (measured as the square root of the Jensen-Shannon distance between the male and female splicing distributions, see methods) was higher in non-SA genes than for SA genes (Figure 7A-C), although this difference was generally small and only significant in larvae (Figure 7A). We also found that genes with a high intersexual genetic correlation tended to exhibit lower levels of differential splicing between the sexes (Figure 7D). The negative correlation between r_mf_ and √JS_(m,f)_ was significant for larvae (r^2^=-0.070, P=0.007) and adults (r^2^=-0.081, P<0.001) but not for pupae (r^2^=-0.038, P=0.110).

**Figure 7.**
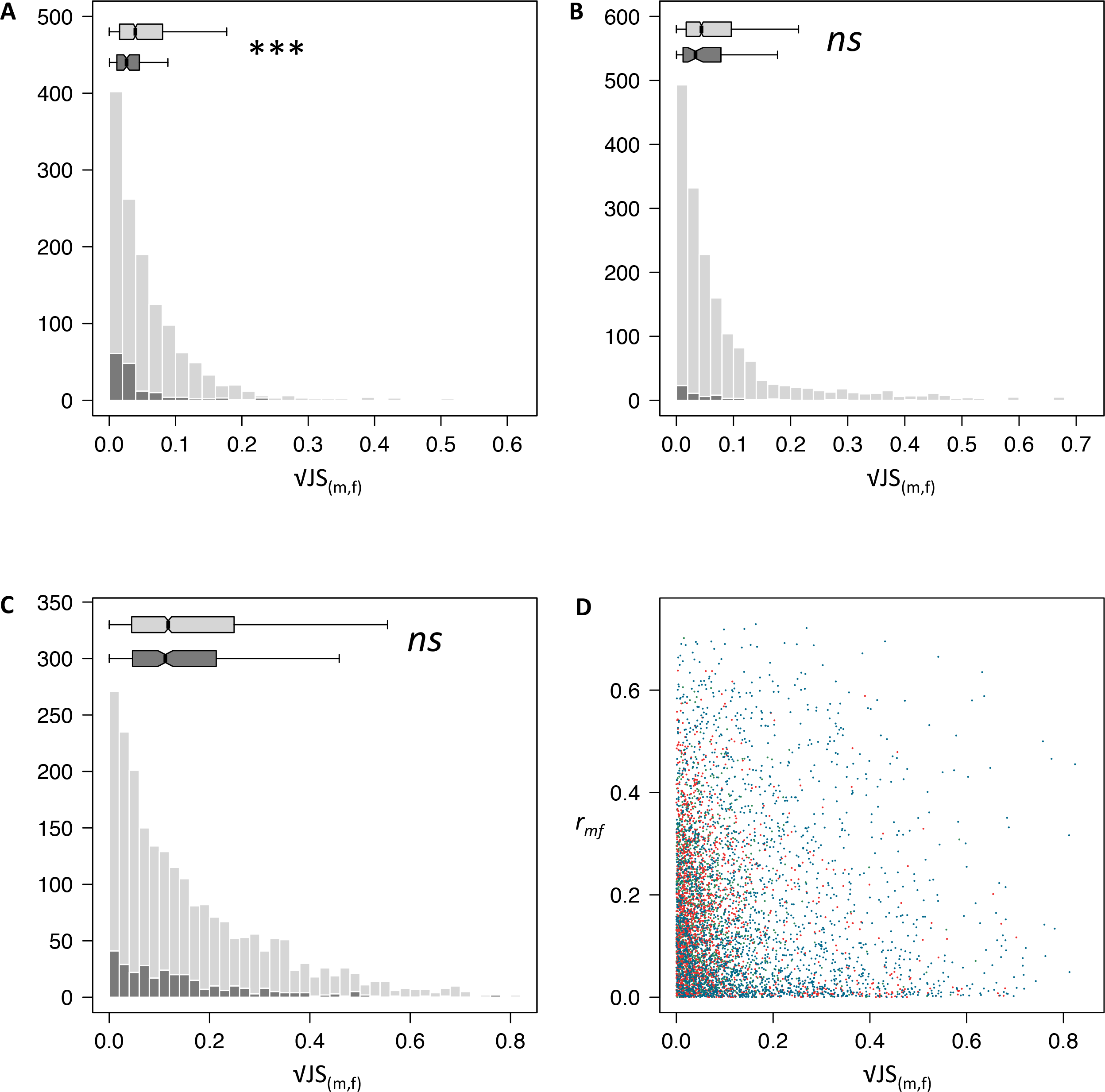
Evidence for differential splicing between males and females in (A) larvae (1636 genes tested); (B) pupae (2015 genes); and (C) adults (2822 genes), as histograms of the square root of the Jensen-Shannon distance. Sexually antagonistic genes are shown in dark grey; non-sexually antagonistic genes in pale grey. The numbers of sexually antagonistic genes tested was generally low (151,63 and 306 genes respectively for larvae, pupae and adults). Notches in boxplots represent ±1.58*IQR/√N. Asterisks indicate significance of the difference between SA/non-SA genes at each stage. (D) The square root of the Jensen-Shannon distance plotted against the intersexual genetic correlation (r_mf_) for larvae (green points), pupae (red) and adults (blue).

## Discussion

Our results clearly demonstrate that although the developmental transcriptome of *D. melanogaster* exhibits high levels of sexual dimorphism, sexual conflict over the expression of shared genes persists. In fact, 18.4% of the genes tested were identified as sexually antagonistic candidates in at least one stage of development, implying that this dimorphism is not a signature of fully resolved conflict. Furthermore, our analyses provide a detailed account of how conflict could be mediated at the level of the transcriptome, as we find evidence for a number of different sources of constraint that could prevent conflict resolution, as well as finding evidence to support one specific mechanism of resolution.

First, the overall patterns of sex-specific selection and sex-specific genetic variation - the two main ingredients necessary for independent trait evolution between the sexes^25^ - vary considerably throughout development. Here, we used lifetime reproductive success (LRS) to estimate selection on genes at each stage. From an evolutionary perspective, LRS provides arguably the most relevant approximation of fitness: selection will favour individuals who leave behind the most offspring (i.e. have the highest LRS). If there is a significant correlation between LRS and the pre-adult expression of a particular gene, then it suggests that the gene might contribute to an aspect of development that affects LRS. Sex- specific selection on gene expression appears inconsistent from one developmental stage to the next, with very little overlap between genes that were identified as either SA or SC across stages. This is perhaps unsurprising given previous research that has demonstrated changes in sex-specific selection through development using different experimental approaches^9-26^. The numbers of genes with significant sex-specific genetic variation also varied across development, and were especially low in larvae and, to some extent, adults, limiting the potential for independent evolution between the sexes at these stages.

Second, the results revealed potential genetic constraints on conflict resolution that stemmed from several sources of shared genetic variation: between-gene, between-sex and between-stage, and sometimes a combination of these. Even if there is considerable overall genetic variation for gene expression, if this genetic variation is not independent between contexts where the gene is under different selection, then adaptive evolution of gene expression can be constrained. It is unlikely that any gene would be expressed completely independently^19-27^; however, the extent of genetic covariance is almost always significantly higher for SA genes than it is for SC genes, strongly supporting the idea that this shared genetic variation could prevent conflict resolution. The exception to this was that gene connectivity was significantly lower for pupal SA genes than for pupal SC genes, and similarly, between-sex and between-gene multivariate shared genetic variation in the B matrix was lower for pupal SA than SC genes. This may result from the unusual modularity of the pupal transcriptome, where most fitness-related genes clustered into a single large, highly correlated gene module that consisted predominantly of SC fitness-related genes.

Consistent with previous research^28^, univariate between-sex genetic covariance and the intersexual genetic correlation is higher for SA genes than for non-SA genes in our data, indicating a putative constraint of genetic variation shared between the sexes within developmental stages. However, the multivariate analyses in this study provide additional insight into between-sex and between-gene genetic covariance that has previously been overlooked. Genetic covariance within the **B** matrix seems particularly influential in adults, where the constraint imposed by **B** on the independent evolution of SA genes between the sexes is high. The results highlight the instability of the genetic covariance in **B** across development, and emphasise the usefulness of a multivariate perspective to examine trait evolution in a more realistic, multi-gene context.

Of particular interest was the amount of shared genetic variation between both sex and stage. This genetic covariance indicates that the expression of a gene in, for example, male larvae, is not independent of the expression of the same gene in, for example, adult females. Since the fitness consequences for a particular gene are unlikely to be aligned between two different sexes and developmental stages, this could be a source of genetic constraint. This genetic covariance has not, to our knowledge, previously been measured for gene expression, but it might be expected that such covariance would be relatively low and unimportant due to the intuitively weak link between different sexes and stages. In fact, this covariance is of a similar magnitude to the other sources of genetic covariance that were measured, and, also similarly to the other genetic covariance components, it is significantly higher in SA than non-SA genes. This could have implications for antagonistic pleiotropy between developmental stages^6-29-30^, suggesting that such relationships might be sex-specific. We also find higher genetic covariance identified for SA genes between stages within each sex than for non-SA genes, supporting a link between developmental antagonistic pleiotropy and sexual conflict^6^. Given the magnitude of between-stage, and between-sex/between-stage, genetic covariance measured here, developmental genetics could have important consequences for conflict resolution. Such developmental covariance has been studied outside of the sexual conflict literature^31^, but this is clearly also relevant to sexual conflict and the evolution of sexual dimorphism.

Finally, we present some limited evidence to support the idea that a lack of differential splicing between sexes might also hinder conflict resolution. Previous research has considered a role of sex-specific splicing^32-33^ or differential exon usage and duplication^34^ in allowing the sexes to achieve differential expression from a shared gene. Consistent with this, we find firstly that larval SA genes have significantly less evidence of sex-specific splicing patterns than non-SA genes, and secondly that the intersexual genetic correlation and the extent of sex-specific splicing are significantly negatively associated in larvae and adults - i.e. genes with more shared quantitative genetic variation between the sexes also exhibit less evidence of sex-specific splicing patterns. This indicates another potential genetic constraint on the independent expression of shared genes between the sexes.

We have therefore identified multiple routes through which conflict resolution could be constrained or facilitated. The novel developmental perspective of this quantitative genomic data is particularly interesting not only because it allowed us to identify genetic covariance between stages, but also because it highlights the different dynamics of sexual conflict at each stage, and the potential to underestimate the extent of conflict or resolution by focussing on only one stage. In short, the larval and adult stages appear to be characterised by conflict and constraint on sex-specific phenotypes, whereas there is less evidence of this in pupae. This is attributable to a complex combination of constraints on larvae and adults that ultimately result in the transcriptome being less independent between the sexes. Since the pupae undergo dramatic metamorphosis and differentiation, compared to the relatively stable larvae and adults, it seems likely that strong selection on the metamorphic process for the formation of optimised sex-specific phenotypes via tissue differentiation may have lead to more extensive conflict resolution in pupae. Indeed, these results are in line with previous work on *D. melanogaster*, which demonstrated some of the most dynamic patterns of gene expression at the start and end of the pupal stage^8^, indicating that this developmental stage may have evolved a more flexible transcriptome to allow for sex-specific metamorphosis, in contrast to larval and adult stages that are characterised by tissue growth and maintenance, respectively. A holometabolous life cycle allows, to some extent, for a discrete phase of concentrated differentiation between the sexes, with potential for some aspects of development to become uncoupled across metamorphosis^35^. Since this phase is absent in hemimetabolous insects, and in other animals more generally, there is potential for conflict to be more prominent during development in other species. It is clear that constraints on conflict resolution are likely to result from a combination of different sources, making the evolution of conflict resolution a complex problem, not least one that appears unlikely to be consistent even throughout the life cycle.

## Methods

Hemiclonal haplotypes were sampled from a *D. melanogaster* base population (LH_M_) that had been maintained in the laboratory for more than 500 generations as a large outbred population with overlapping generations. Haplotypes were expressed as male or female hemiclonal individuals following a series of crosses^17^’^26^. All flies were reared on a standard molasses diet at 25°C and 65% relative humidity, with a 12:12h light:dark incubator cycle.

Male and female flies used for the parental cross were allowed to interact and mate for 48h before the males were removed and females were flipped into lightly yeasted laying vials. Females oviposited in these vials for 2h before being flipped into a holding vial. Females were given further 2h laying periods in fresh vials after 4 and 7 days, creating staged vials of developing offspring. For each set of vials, larvae were visually inspected under a dissecting microscope after 4 days, when developing testes can be seen through the larval body wall. Larvae were split into sex-specific vials to continue development, with 10 larvae per vial. Eleven days after the initial laying vials were set up, third instar larvae, pupae and 1-day old virgin adults (unable to mate as they eclosed in sex-specific vials) were harvested. Individuals were frozen at -80°C and sample processing took place over the course of 4 weeks.

RNA extractions were carried out on individual larvae/pupae/adults using TRIzol (Invitrogen) according to the manufacturer’s instructions (adjusted protocol for a small amount of starting material). Our aim was to sequence RNA from 180 samples in total, comprising males and females from three developmental stages from each of 10 hemiclonal lines, with three biological replicates per sample type. However, the hemiclone cross produces siblings that do not have the hemiclonal genotype, which are identified as adults with *bw eye* colour among wild-type hemiclonal flies. We therefore carried out additional identification steps for larval and pupal samples, and collected more than twice the number of samples required in order to compensate for non-hemiclonal genotypes. After the initial RNA extraction process (but prior to DNase treatment), a subsample of the extract from larvae/pupae was used in PCR reactions to identify the correct genotypes. Two PCR reactions were carried out: one set of primers was designed for the dominant insertion mutation in the *bw gene, bw^D^* (F: CTTATCTTTGGAGAGAAGAGA; R: GGATCATCCGTGCATCAAGAC), and the other set for the male fertility factor *kl5* on the Y chromosome (F: GCTGCCGAGCGACAGAAAATAATGACT; R: GTCCCAGTTACGGTTCGGGTTCCATTGT), to make sure the sexes had been identified correctly by phenotype. Primers to a region on chromosome 2R were used as a control reaction (F: AAAAGGTACCCGCAATATAACCCAATAATTT; R: GTCCCAGTTACGGTTCGGGTTCCATTGT). Control and Y chromosome primer sequences were taken from Lott et al.^36^.

After the genotypes had been identified, DNase digestion was carried out on the correct samples, and RNA was suspended in RNase-free water. Library preparation (Illumina TruSeq RiboZero) and Illumina HiSeq sequencing was carried out by AROS Applied Biotechnology (Aarhus, Denmark) according to Illumina v.4 protocols, using 30 sequencing lanes with 6 samples per lane. For each sample, 400ng of total RNA was used and 32-49M reads were achieved per sample. Gene expression data were aligned to the FlyBase version r6.05 of the *D. melanogaster* genome^37^ and normalised within the R (v3.2.1) BioConductor ‘QuasR’ and ‘DESeq’ pipeline^38^. At this stage, samples were checked for the *bw* gene and the fertility factor *kl5*, and one sample was removed due to having been incorrectly genotyped prior to sequencing (a male pupa had been incorrectly assigned as a female pupa from one of the hemiclonal lines), leaving 179 samples for analysis. The alignment generated read counts for 16727 genes. The dataset was filtered to remove low variance genes, leaving 14008 genes, of which 13501, 13602 and 13495 were expressed in larvae, pupae and adults respectively.

Sex-specific fitness data was recorded as hemiclonal line averages of male and female total reproductive success under competitive conditions, as described in assays in previous studies^17-26^. Fitness data was analysed as relative fitness by dividing line-specific fitness means by the maximum average fitness for each sex.

**Sex-biased and fitness-associated gene expression**. Variation in the read counts for each gene was partitioned in a generalised linear mixed model using the ‘glmer’ function in the R package ‘lme4’^39^ across sex, line and sex-by-line terms. Initially, a separate model was performed for each gene at each developmental stage. This was due to the difficulties in model convergence trying to run a full model (which would require sex-by-line-by-stage-by-fitness effects) with data from only 10 genetic lines. Models assumed a Poisson distribution and included ID as a random term to account for differences in total number of reads per sample. The significance of variance components was calculated from a 0.5X^02^+0.5X_1_^2^ mixture distribution from a likelihood ratio test comparing the full model with the reduced model (without the component being tested), and *P* values were adjusted according to the false discovery rate (FDR)^40^ including the full set of results from all three developmental stages (since most genes were tested at every stage). We then tested if each gene was associated with fitness at each stage, using a GLMM under the same conditions as before but with sex, fitness and fitness*sex. A significant fitness*sex interaction was interpreted as sexually antagonistic (SA) selection, whereas a significant overall fitness term, with a non-significant fitness*sex interaction, was interpreted as sexually concordant (SC) selection.

**Transcription modules**. We began by calculating the genetic correlation between all pairwise combinations of fitness-related (both SA and SC combined) genes separately in larvae (N=1261), pupae (N=3103) and adults (N=2959). The absolute correlations were used to examine overall gene connectivity (average of all absolute pairwise correlations per gene at each stage), and were then used in a clustering algorithm to identify gene modules^21^. This algorithm works via an optimisation function that finds a pattern of gene modules that maximises correlation within modules while minimising correlation between modules. Each of the identified modules was tested for enrichment with SA genes using hypergeometric tests (P values adjusted for FDR). Next, we identified tissue- specific genes for larval and adult tissues using the FlyAtlas *D. melanogaster* microarray database^22^, and tested modules for enrichment with tissue-specific genes. These tests could only be carried out on modules with >100 genes due to sample size issues. Again, hypergeometric tests were used and P values were FDR-adjusted.

**Quantitative genetic analyses**. In order to estimate the full univariate between-sex and between-stage genetic variance-covariance matrix for each gene, we used the ‘MCMCglmm’ package^41^ in R to partition variance in gene expression for the 14008 genes that varied in expression at any stage. The posterior distribution for each model was estimated using weakly informative inverse-Wishart priors for the variance components (V=diag(6)/6, nu=6) and convergence was visually inspected with diagnostic plots. The models included stage and sex as fixed terms, and partitioned genetic variance across a 6x6 matrix for each gene, with female- and male-specific larval, pupal and adult genetic variance and all possible covariances. The posterior distribution of these models was used for all univariate quantitative genetic aspects of the analysis. We calculated the intersexual genetic correlation, r_mf_, from the posterior estimates of variance components^42^; and used the absolute sum of the relevant variance components (scaled to the total amount of genetic variation in the full matrix) to compare the amount of between-sex and/or between-stage shared genetic variation across SA and non-SA genes.

We also examined shared genetic variation between-sex and between-gene for each developmental stage independently. Ideally, this would involve running a multivariate model with expression variance for all genes partitioned between sex and line. However, this would involve estimating a genetic variance-covariance matrix (**G** matrix) with more than 5 million parameters. Instead, we ran 2000 iterations of a multivariate model that randomly sampled 5 genes at a time, and, using the expression of the 5 genes as a multivariate response, partitioned gene expression variance between sex and line. Each model was ran using ‘MCMCglmm’^41^ in R, with sex:gene as fixed terms, sex:gene:line as random terms, and sex- and gene-specific residuals. The prior specified a weakly informative inverse-Wishart distribution for the variance components (V=diag(10)/10, nu=10), and model checks were carried out as above. Each iteration of the model generated a posterior distribution of a 10x10 **G** matrix (5 genes expressed in 2 sexes). We repeated the analysis for six subsets of genes, which were exclusive within each developmental stage: SA and SC genes in larvae, pupae and adults. Results are summarised from the average **G** matrix calculated from the posterior distribution of all 2000 model iterations. Importantly, this **G** matrix represents the full Gmfmatrix, including the submatrix **B**, which summarises the between-sex and between-gene covariance (see Supplmentary File 3 for details). We examined genetic variation within **∥B∥** using two metrics. First, **∥B∥**, the matrix norm was calculated to approximate the magnitude of the variation within **B**. The matrix norm was used instead of the matrix trace (that has been used previously to estimate the amount of variance in **G**) since the **B** sub-matrix is composed entirely of covariance estimates and can therefore have negative eigenvalues on the main diagonal. Note, however, that using the sum of the absolute eigenvalues of **B** (i.e. the **B** matrix trace) produces qualitatively identical results. Second, the matrix correlation between the upper and lower halves of **B** was calculated to estimate symmetry of variance within **B**. Finally, we implemented the multivariate breeder’s equation: 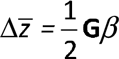(Lande 1980). This equation uses the product of the genetic variation in **G** and the vector of linear selection gradients in *β* to calculate a vector of the predicted response to selection for each trait, 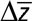. We generated a vector of linear selection gradients that corresponded to each estimate of the **G** matrix in the models described above. For each iteration of the **G** matrix model, selection gradients for the same 5 genes were estimated from a multiple linear regression of the genes against relative fitness for males and females, following Lande^24^ This allowed us to calculate sex-specific 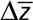 based on the sex-specific components of **G** and *β*. This calculation was carried out using the distribution of 2000 estimates, to create a distribution of 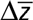 estimates, from which confidence intervals could be used to assess significant differences.

Finally, to include B in the calculation of 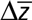, the multivariate breeder’s equation^24^ can be expanded as:

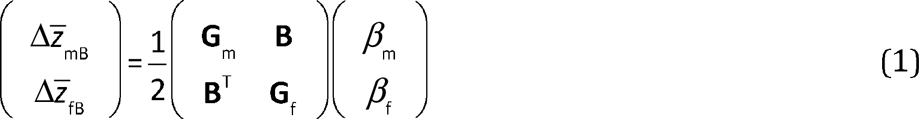

where 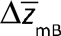 and 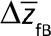 represent the predicted response to selection of each sex given both the sex-specific genetic variation in **G_m_** and **G_f_** and the shared genetic variation within **B**. The predicted divergence between the male and female response to selection was measured as the angle between male and female 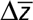with and without **B**. The angle was calculated as:

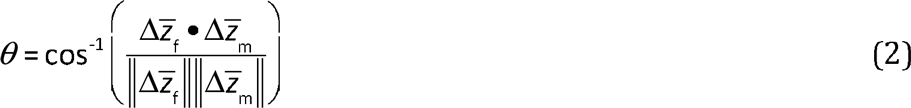

with 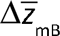 and 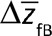 substituted for the calculations accounting for shared genetic variation in **B**. As before, all calculations used the distribution of 2000 estimates from the original models to enable confidence interval calculation and significance testing.

**Differential splicing**. We used the cuffdiff function in Cufflinks software^43^ to test for evidence of differential splicing between males and females at each stage. This test runs for any gene where alternative isoform expression is found, which in this dataset tested 1636, 2015 and 2822 genes in larvae, pupae and adults respectively. The analysis uses the Jensen-Shannon distribution to test for significant differential splicing between the male and female samples within each developmental stage.

Accession codes. RNA-seq data have been deposited with accession SRP068235

## Acknowledgments

This work was supported by grants from the Swedish Research Council (2011-3701), European Research Council (#280632), and a Royal Society University Research Fellowship (all to E.H.M). The authors thank M. Reuter for his comments on an earlier draft of the manuscript.

## Author Contributions

*F.C.I. and E.H.M. wrote the paper and designed the experiments. T.M.P. and I.F. set up the hemiclone lines and collected fitness data. F.C.I. and C.L.W. collected the samples and performed the molecular work. F.C.I. performed the statistical analyses*.

---

**Supplementary File 1**. List of Entrez Gene IDs tested at each developmental stage, annotated with results of significance tests for fitness association, sexbiased expression and genetic variation.

**Supplementary File 2**. Results from analysis of gene clusters. Gene modules and connectivity of genes within modules are given for each fitness-associated (SA or SC) gene at each developmental stage (identified by Entrez Gene IDs). Results of hypergeometric tests (adjusted for FDR<0.05) for cluster enrichment with (A) SA genes and (B) tissue-specific genes are also provided.

**Supplmentary File** 3. Summary of the structure of the full **G** matrix, **G_m_f**, and how it was used in the multivariate analyses.

